# Engineering antigenic breadth against SARS-CoV-2 by pairing divergent RBDs within a single mRNA immunogen

**DOI:** 10.64898/2026.01.12.699133

**Authors:** Isabelle Montgomerie, Rebecca E. McKenzie, Olga R Palmer, Ngarangi C Mason, Joanna Kuang, Theresa E. Pankhurst, Sarah L. Draper, Sventja von Daake, David A. Eccles, Thomas W. Bird, Abby L. Martin, Isaac Green, Lydia White, Jordan J Minnell, Sam J. Small, Ian F Hermans, Gavin F Painter, James E Ussher, Miguel E. Quiñones-Mateu, Wayne M Patrick, Davide Comoletti, Lisa M. Connor

## Abstract

Vaccines capable of eliciting broadly neutralising antibodies (bnAbs) are a major goal for pandemic preparedness. A persistent challenge across vaccine fields is how to deliberately recruit the rare B cell clones that recognise conserved epitopes shared across diverse viral variants. BnAbs have been known to frequently emerge through extensive somatic hypermutation during affinity maturation, here we describe an alternative, structure-driven mechanism for bnAb selection. We designed an mRNA vaccine in which two antigenically distinct SARS-CoV-2 variant’s (Omicron and Delta; O-Δ) receptor binding domains (RBDs) are physically fused on a single polypeptide. This design is predicted to favour B cell antigen receptors capable of engaging conserved epitopes on both RBDs with enhanced avidity. A matched non-divergent tandem RBD (Delta-Delta; Δ-Δ) served as a control. The divergent (O-Δ) immunogen was robustly expressed and retained high-affinity ACE2 binding. In mice, immunisation elicited potent antibody responses and increased the frequency of antigen-specific cross-reactive B cells, recognising Delta, Omicron, and the 2002 pandemic strain SARS-CoV RBDs. Using multicolour RBD tetramers and single-cell B cell receptor sequencing, we show that breadth arises via two distinct pathways. The divergent vaccine preferentially enriches clonally distinct cross-reactive B cells (not present within non-cross-reactive B cell pools) with low levels of somatic hypermutation (SHM), consistent with selection of germline-biased precursors. In contrast, the matched control vaccine yields cross-reactivity primarily within existing clonal lineages (clonal overlap between cross-reactive and non-cross-reactive cells) and at higher mutational burdens, consistent with affinity-maturation-driven acquisition of breadth. Together, these findings demonstrate that antigen structure can bias B cell selection towards cross-reactive specificities without requiring extensive SHM. This work establishes a simple, modular antigen-design principle in which juxtaposing appropriately divergent antigens on a single scaffold promotes the enrichment of bnAb-prone B cells, providing a scalable strategy for vaccine development against rapidly evolving pathogens.

## Introduction

SARS-CoV-2 has diversified markedly since the late-2019 zoonosis, accumulating immune evasion substitutions, increasing infectivity, and undergoing antigenic drift under global circulation. First-generation vaccines protect well against severe disease but afford limited breadth. Durable, broad-spectrum immunity is therefore a priority for pandemic preparedness. Although infection and current vaccines can generate neutralising antibodies to conserved regions of the spike receptor binding domain (RBD)^1-6^, truly broadly neutralising antibodies (bnAbs) are rare, in the order of ∼1% of neutralising antibodies^5^, indicating a selection bottleneck rather than an absolute feasibility barrier. The challenge lies in the amplification of rare clones that target conserved regions within the RBD. To achieve breadth, responses must focus on functionally constrained, accessible RBD epitopes, such as sites with low mutational tolerance, where substitutions reduce ACE-2 engagement, spike stability, or viral entry^5,7^. In practice, these include compact patches within the ACE2-contact core where bnAbs bind by receptor mimicry^5^ and conserved non-receptor binding motif “silent-face” surfaces (CR3022/site-IV)^8^ that buttress RBD core stability and regulate spike conformational dynamics^5,9^.

Multiple strategies seek to broaden protection against sarbecoviruses (such as SARS-CoV-2) and variants including variant-adapted boosters^10^, chimeric spike constructs^11^, consensus sequence antigens^12,13^, mosaic displays^14-18^, and epitope-focused immunogens^19-21^. Yet many require co-delivery of large or multiple proteins, increasing complexity and manufacturing burden, and can inadvertently reinforce immunodominance (e.g., against non-neutralising regions distal from the receptor binding site, or mutable receptor binding motif subregions)^22-25^ by amplifying high-frequency but escape-prone specificities^26^. Several antibodies once labelled "broadly neutralising" have also lost activity against recently emerged variants^27-29^, demonstrating that breadth is fragile unless anchored to functionally constrained epitopes^5^.

Here we present a new vaccine design strategy to selectively amplify cross reactive clones, by preferentially increasing the copy number of cross protective epitopes. Antibodies bind most effectively when they encounter repeated copies of the same epitope, because repetition enables higher-avidity engagement through multivalent binding. We therefore aimed to increase the local copy number of conserved epitopes by pairing two divergent RBDs in nanoscale proximity within the same molecule. Pairing antigenically divergent RBDs ensures that only conserved epitopes appear in repeated form, because variable (strain-specific) epitopes differ between the two RBDs and therefore lose the multivalent advantage. In contrast, in a construct where the RBDs are from matched strains, every epitope is duplicated, so B cells targeting any site, whether conserved or variable receive an equal avidity advantage. This favours the expansion of many high-affinity, strain-specific clones but provides no selective pressure for recognising conserved regions. Therefore, we hypothesised that pairing divergent RBDs would shift the competitive landscape: fewer epitopes support high-avidity binding, but those that do are conserved features most likely to generate cross-reactive antibodies.

To test this hypothesis we engineered a compact, single open reading frame (ORF), mRNA-encoded immunogen. This vaccine was based on our previous protein-based design comprising two identical RBDs and an N terminal domain (NTD) translocated to the C-terminus, in a single polypeptide chain^30^. Our new vaccine was adapted to be delivered as an mRNA vaccine, and a transmembrane domain (TMD) was added such that the protein was expressed on the cell surface. In our new vaccine two antigenically divergent RBDs (SARS-CoV-2 variants: Delta and Omicron) were cis-linked on the cell membrane, while retaining the NTD to supply robust CD4 T cell help via abundant class II epitopes^30,31^. This compact genetic design is smaller than the full-length spike and avoids multi-antigen cocktails or large protein nanoparticles, while preserving the principle of presenting divergent antigens in close proximity to preferentially expand B cell clones with receptors recognising shared (conserved) determinants. This creates a direct molecular route to expanding cross-reactive clones and provides a fundamentally new framework for designing broadly protective vaccines. We compare this new divergent construct to a “matched” construct, in which both RBDs within the polypeptide were derived from the Delta variant.

Using multicolour flow cytometry and fluorescent RBD tetramers (B cell antigen baits), we show that this tandem-RBD construct increases the frequency and breadth of antigen-specific B cells and, importantly, also enriches for cross-reactive clones. These clones are phylogenetically distinct from non-cross-reactive clones and carry a relatively low mutational load, consistent with germline-biased selection of conserved-epitope B cells rather than by prolonged maturation. These data shown that a single-molecule, mRNA-encoded geometry underpins a structural paradigm for amplifying cross-reactive humoral immunity and offer a new design principle for broadly protective SARS-CoV-2 vaccine development.

## Results

### Membrane-anchored tandem RBD is robustly expressed and ACE2-accessible

We developed an mRNA version of our previously described RBD-RBD-NTD protein immunogen^30^ to directly test whether antigen geometry and epitope repetition can bias germinal centre competition toward B cells recognising conserved RBD surfaces. We converted the design into a compact, single-ORF, membrane-anchored immunogen by adding the spike transmembrane (TM) domain yielding a tandem RBD + NTD construct (tRBD-NTD) in which two RBDs and one NTD are encoded in a single polypeptide (Fig. 1a, b). To isolate the effect of inter-variant divergence while holding antigen format constant, we generated two pairing types: a divergent tRBD-NTD with RBDs from two different SARS-CoV-2 variants, Delta and Omicron BA.2 (O-Δ); and a matched (non-divergent) control tRBD-NTD containing two Delta RBDs (Δ-Δ) RBDs (Fig. 1b). In the divergent (O-Δ) construct, only conserved surfaces are present in duplicate, whereas strain-specific epitopes are not; this design therefore provides a direct way to test whether duplication of conserved determinants selectively favours cross-reactive B cell clones in vivo.

**Figure 1.**
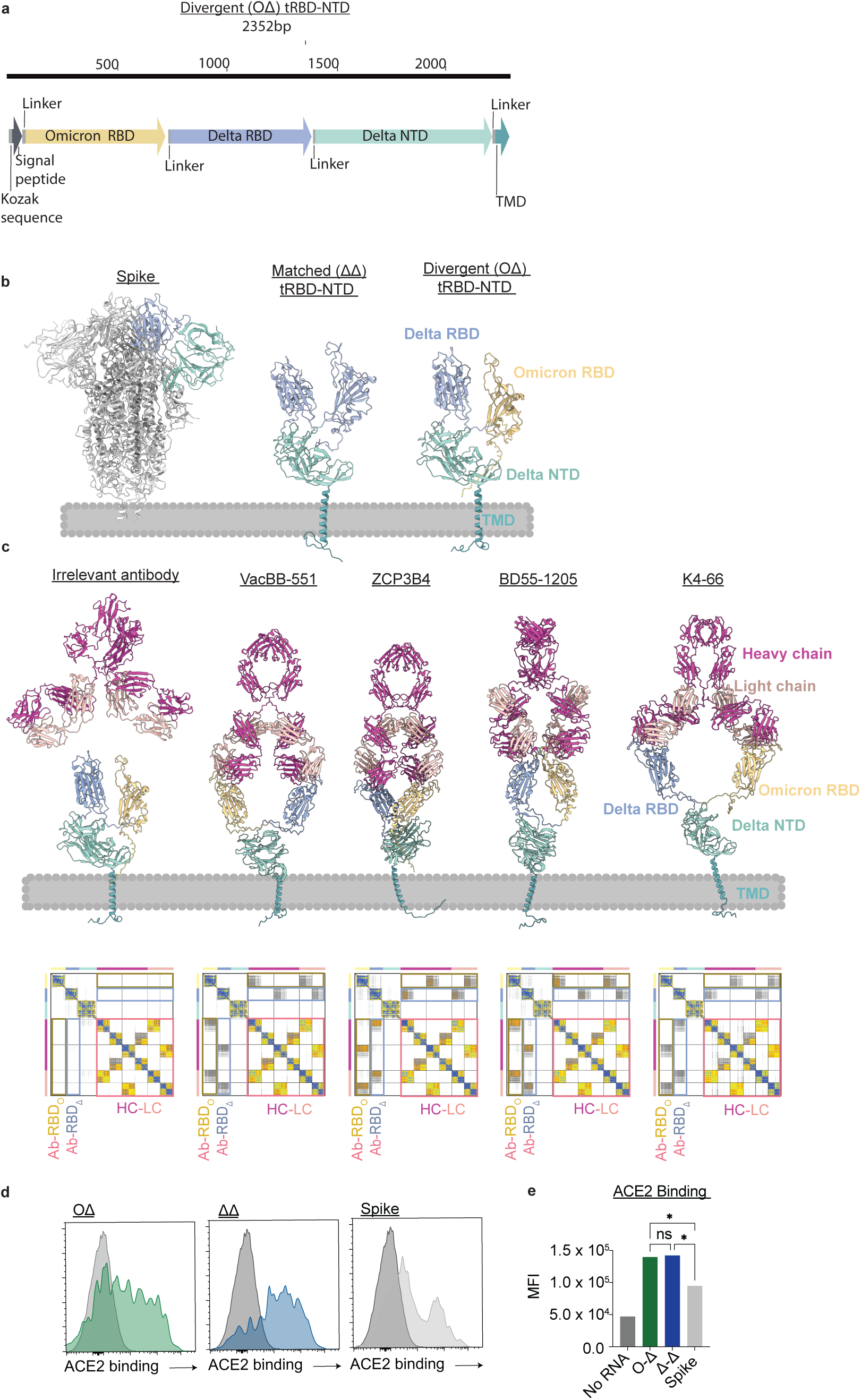
Design and expression of membrane-anchored tandem RBD-NTD (tRBD-NTD) mRNA immunogens. a, Single open reading frame schematic encoding two RBDs and one NTD with a C-terminal transmembrane (TM) domain. Two pairings were tested: matched control (Δ-Δ) and divergent (O-Δ), the Omicron RBD stabilised by removing the following mutations: S371L, S375F, K417N, N501Y. **b,** AlphaFold3 models of tRBD-NTD (Δ-Δ and O-Δ) immunogens. **c,** AlphaFold3 predicted binding of tRBD-NTD (O-Δ) to broadly reactive anti-RBD monoclonals (VacBB-551, ZCP3b4, BD55-1205, and K4-66), illustrating plausible engagement of both RBDs (PAE maps shown). Below models are PAE plots quantifying binding confidence: low PAE (blue) indicates high confidence, high PAE (grey) indicates low confidence. **d,** Surface expression and ACE2 accessibility of immunogens in HEK293T cells transfected with each vaccine construct and full-length spike (positive control) and stained with hACE2-Fc detected by anti-human Fc-PE. **e,** Geometric mean fluorescence intensity (GeoMFI) quantification. Bars represent geometric mean, statistical analysis using one-way ANOVA with a Fisher post-test. P value * is <0.1.

We then asked whether the geometry of tRBD-NTD could plausibly support bivalent engagement of conserved epitopes (intramolecular bivalency) by a single BCR. In silico modelling of O-Δ tRBD-NTD in complex with well-characterised broadly neutralising antibodies targeting conserved ACE2-focused epitopes (VacBB-551^32^, ZCP3B4^33^, BD55-1205^5^, and K4-66^34^), using AlphaFold3, visualised in UCSF ChimeraX (version 1.9), produced confident folds for each RBD and antibody variable domain and low-to-moderate predicted error at the RBD-antibody interfaces for both Delta and Omicron RBDs. (Fig. 1c). An irrelevant antibody with no affinity for SARS-CoV-2^35^ did not yield a confident interface (negative control). While these models are hypothesis-generating rather than structural evidence, they support the feasibility of simultaneous engagement of conserved epitopes on both RBDs.

An additional non-mutually exclusive mechanism is that cross reactive B cells are amplified by intermolecular BCR cross-linking, where repeated conserved sites can cluster multiple BCRs on the same cell and thereby increase avidity, a concept consistent with prior reports showing that duplicating RBD enhances BCR cross-linking and immunogenicity^36,37^, and our original design linked two RBDs explicitly for this purpose^30^.

To enable a fair comparison between matched and divergent tRBD-NTD formats, we ensured equivalent antigen expression. Because Omicron BA.2 RBD can express poorly and included mutations that were associated with reduced expression, as determined by protein expression in HEK293 cells, we reverted four residues (S371L, S375F, K417N, N501Y; amino acid position annotated based on full spike sequence) back to the ancestral Wuhan sequence to improve O-Δ expression to levels comparable to Δ-Δ tRBD-NTD. HEK293T cells transfected with each mRNA construct showed ACE2 binding using Fc-hACE2/anti-human Fc-FITC probe and flow cytometry, indicating correctly folded ACE2-accessible RBD, and robust surface display for both formats relative to full-length spike (Fig. 1d, e), thereby equalising antigen availability for immunisation. Finally, both tRBD-NTD formats were immunogenic in mice. Two intramuscular (i.m.) doses given three weeks apart elicited comparable serum anti-RBD IgG titres and avidity (Fig. 2a-c). In a separate booster cohort primed with ancestral full-length spike and boosted at day 176, both Δ-Δ and O-Δ tRBD-NTD robustly increased RBD-specific titres and sustained elevated levels to day 400 (Fig. 2d-e). Together these results validate the matched/divergent platform and set the stage for the mechanistic dissection of cross-reactivity by B cells.

**Figure 2.**
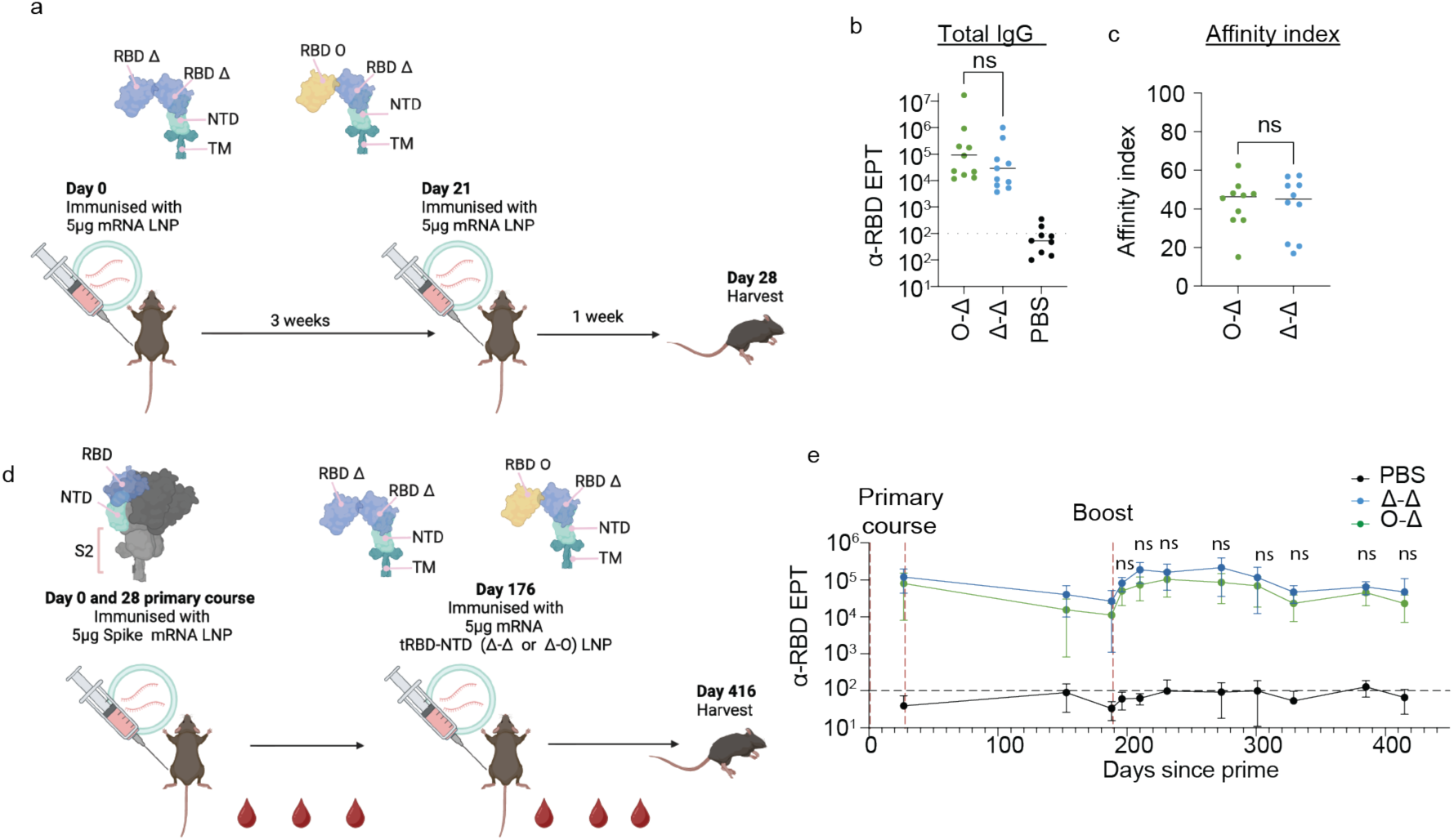
Immunogenicity of matched (Δ-Δ) and divergent (O-Δ) tRBD-NTD mRNA vaccines. a, Immunisation scheme: mice received intramuscular injection of 5 ug mRNA with N1-methylpseudouridine formulated in ALC-0315 containing ionizable lipid nanoparticles twice, three weeks apart and were analysed 7 days after the second shot. **b,** Anti-RBD IgG titres (ELISA) after two doses for full-length spike (benchmark) and tRBD-NTD (Δ-Δ and O-Δ). **c,** Avidity index of anti-RBD IgG. **d,** Booster study design: priming with ancestral full-length spike on days 0 and 21, followed by a tRBD-NTD (Δ-Δ and O-Δ) booster on day 176. **e,** RBD-binding IgG titres following booster immunisation, tracked longitudinally to day 400. Solid bars represent mean, statistical analysis (**b**) using one-way ANOVA with a Sìdák post-test (**c**) Mann-Whitney test, and (**e**) two-way ANOVA with a Fisher post-test. P value * is <0.05.

### Divergent tRBD-NTD vaccine elicits antigen-specific B cells and broadens neutralising responses against SARS-CoV-2 variants without loss of magnitude

To characterise how the matched (Δ-Δ) and divergent (O-Δ) tRBD-NTD vaccines perform at the level of antigen-specific B cells and functional antibodies, we first evaluated vaccine-specific B cell repertoires using multicolour RBD tetramers for Delta, Omicron (BA5) and SARS-CoV (2002 pandemic strain, hereafter referred to as CoV-1). Mice were immunised twice three weeks apart (Fig. 2a) and antigen specific B cells in the draining lymph node (inguinal lymph node) were analysed. Flow cytometry revealed substantial RBD-binding B cell populations for all three tetramers in both vaccine groups (Fig. 3a-c). Notably, the divergent (O-Δ) tRBD-NTD yielded higher frequencies across Delta, Omicron and CoV-1-binding B cells than the matched (Δ-Δ) tRBD-NTD (Fig. 3b-d), indicating a broader expansion of antigen specific B cells that includes recognition of a more distant sarbecovirus. This result demonstrates that the divergent construct increases the breadth of the B cell response at the population level.

**Figure 3.**
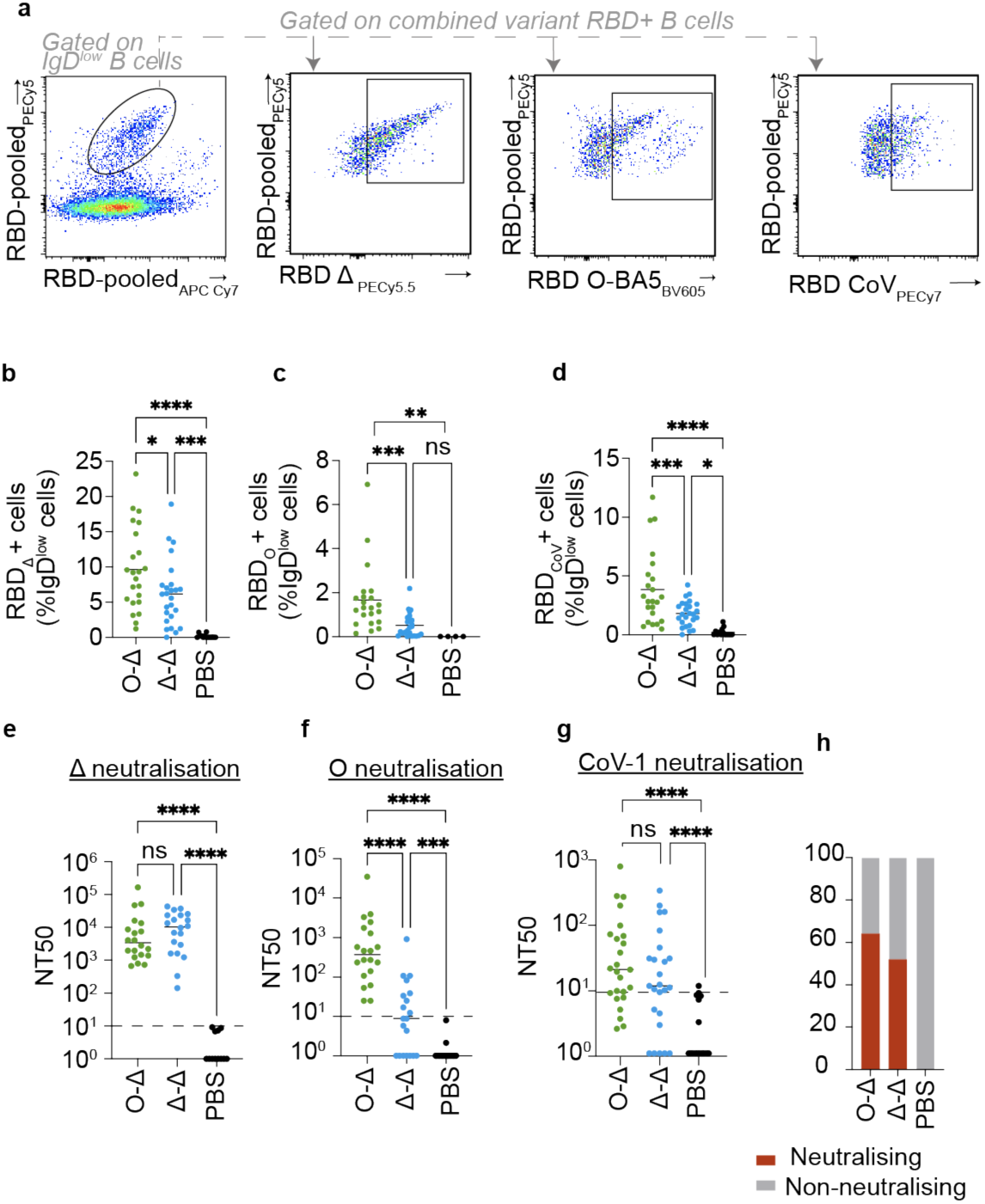
Divergent (O-Δ) tRBD-NTD vaccine induces antigen-specific B cells and neutralising antibodies. a-h Mice were immunised twice, 3 weeks apart, and analysed 7 days after the second shot, draining lymph nodes (dLN) were harvested for antigen specific B cell analysis and serum for neutralisation assays. RBD proteins (Delta, Omicron BA.5 and SARS-CoV-1) were enzymatically biotinylated with BirA and assembled into tetramers with fluorescent streptavidins. To capture the total RBD-specific pool, each RBD tetramer was prepared in PE-Cy7 and APC-Cy7 forms; in addition, each RBD tetramer carried a unique detection fluorophore to permit unambiguous discrimination of strain-specific and cross-binding B cells. **a,** Representative flow plots of B220+, IgD^low^ RBD binding B cells (irrelevant bait-excluded), in dLNs (**b-d**), Frequencies of RBD binding B cells quantified with tetramers for Delta (b), Omicron (**c**), and SARS-CoV-1 (**d**). **e–g**, Serum pseudovirus neutralisation against Delta (**e**), Omicron (**f**), and SARS-CoV-1 (**g**) spikes; horizontal dotted line denotes the assay detection threshold. **h**, Proportion of mice with detectable SARS-CoV-1 neutralisation (NT_50_ above baseline). Data are representative of three or more independent experiments. Cross bars represent mean, statistical analysis using one-way ANOVA with a Fisher post-test. P value * is <0.05, ** is <0.01, *** is <0.001, **** is <0.0001.

We next assessed neutralising activity using pseudovirus neutralisation assays. Both vaccines generated neutralising antibodies against Delta, with matched tRBD-NTD vaccine showing higher Delta NT50 titres (Fig. 3d) despite having a smaller Delta-binding B cell-pool, a finding that suggests qualitative differences can decouple neutralisation potency from the size of the strain-specific B cell pool. As expected from antigen-composition, divergent tRBD-NTD vaccine elicited stronger Omicron neutralisation and a larger Omicron-specific B-cell compartment (Fig. 3c, f). CoV-1 neutralisation was low reflecting antigenic distance, but a greater fraction of O-Δ-vaccinated animals exceeding the assay threshold (NT_50_ above unvaccinated baseline) compared with Δ-Δ (Fig 3g, h). Together, these results reveal a fundamental distinction: the O-Δ design expands cross-recognition across Delta, Omicron and a distant sarbecovirus, and this breadth occurs without wholesale loss of potency, maintaining overall immunogenicity, while matched tRBD-NTD sharpens strain-specific neutralisation. In short, divergent tRBD-NTD vaccine gains breadth by actively redirecting immunity toward broader B cell recognition and cross-neutralisation, without compromising magnitude.

### Single-cell RBD specificity mapping reveals that the divergent (O-Δ) tRBD-NTD expands both variant-specific and cross-reactive B cells

An unresolved question in vaccinology is whether optimal antibody breadth is produced by recruiting multiple variant-specific B cell lineages or by selectively expanding genuinely cross-reactive clones that recognise conserved epitopes. Our divergent (O-Δ) tRBD-NTD vaccine is uniquely suited to test this: it can in principle recruit variant-specific responses (because it contains Omicron and Delta RBDs) but, crucially, duplicates only conserved determinants, which should preferentially favour B cells that bind shared sites if the avidity hypothesis holds. To distinguish these alternatives, we profiled single-cell antigen binding using multicolour RBD tetramers for Delta, Omicron (Fig. 4a-b), CoV-1 (Fig. 4b-c), and ancestral Wuhan-Hu-1 (Fig. 4e-f) on draining LN B cells one week after boost (Fig. 2a). Multicolour flow cytometry allowed us to quantify co-binding of individual B cells to defined B-cell-tetramer combinations and thereby distinguish variant-specific cells (e.g. Omicron-only) from B cells that recognise shared epitopes across variants (e.g. B cells that bind both Omicron and Delta RBD tetramers).

**Figure 4.**
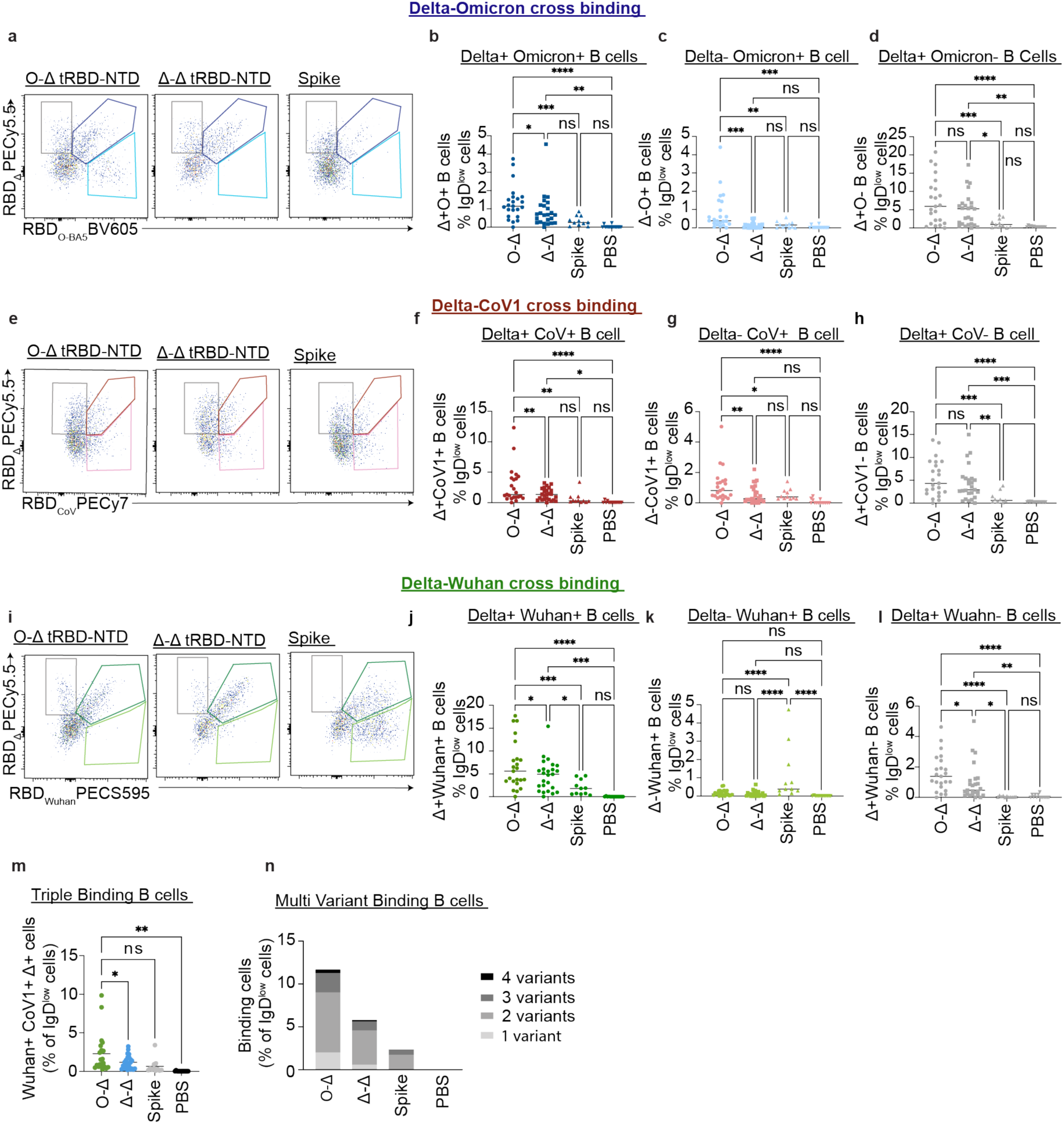
Divergent (O-Δ) tRBD-NTD expands broadly cross-reactive B cells that bind multiple sarbecovirus RBDs. a-l, Mice were immunised twice, 3 weeks apart and inguinal lymph nodes were analysed 7 days after the second shot. B220+ IgD^low^ B cells were profiled with multicolour RBD tetramer baits. **a-d**, Delta and Omicron binding. (a) Representative gating for Delta+ Omicron+ co-binders. Frequencies of Omicron+ Delta+ **(b)** Omicron+ Delta- **(c),** and Delta+ Omicron- **(d)** B cells**. e-h,** Delta and CoV-1 binding. (e) Representative gating for Delta+ CoV-1+ co-binders. Frequencies of Delta+ CoV-1+ **(f)** CoV-1+ Delta- **(g)** and Delta+ CoV-1- **(h)** B cells. **i-l**, Delta and Wuhan (ancestral) binding. (**i**) Representative gating for Delta and Wuhan co-binders. Frequency of Wuhan+ Delta+ **(j)** Wuhan+ Delta- **(k)** and Delta+ Wuhan- **(l)** B cells**. m-n,** Breadth of B cell reactivity across four RBD variants (Delta, Omicron, Wuhan, SARS-CoV-1). (**m**) Frequency of B cells binding Delta, CoV1, and Wuhan. (**n**) Distribution of antigen-specific B cells binding 1, 2, 3, or 4 RBDs. Data are representative of three or more independent experiments. Cross bars represent mean, statistical analysis using one-way ANOVA with a Fischer post-test. P value * is <0.05, ** is <0.01, *** is <0.001, **** is <0.0001.

The divergent (O-Δ) tRBD-NTD vaccination increases the frequency of multi-binding B cells rather than merely adding more variant-specific lineages. Specifically, while the divergent (O-Δ) vaccine induced a large Omicron-specific B cell population relative to the matched (Δ-Δ) tRBD-NTD control (Fig. 4a-c), it also drove a significantly higher frequency of Delta+ Omicron+ double-binding B cells (Fig. 4a-b), whereas the frequency of Delta+ Omicron-cells was similar between the matched (Δ-Δ) and divergent (O-Δ) tRBD-NTD immunised mice. Thus, the divergent construct does more than introduce Omicron-only specificity, it selectively expands cross-reactive clones capable of binding both RBDs in the vaccine.

To evaluate recognition of conserved surfaces independent of Omicron inclusion, we quantified co-binding to Delta and CoV-1 tetramers. Although neither vaccine encodes CoV-1 antigen, divergent (O-Δ) immunisation increased the frequency of Delta+ CoV-1+ double binding B cells and, notably, revealed a small CoV-1-only population, that did not bind the vaccine-encoded Delta (Fig. 4e-h). A similar enrichment was observed for Delta and Wuhan, Delta+ Wuhan+ co-binding B cells were higher after divergent (O-Δ) tRBD-NTD immunisation (Fig. 4i-l). Finally, Boolean gating across all tetramers showed that both the number and proportion of multi-binding B cells (>3 tetramers) were greater after divergent (O-Δ) tRBD-NTD vaccination, alongside an expansion of the total antigen-specific pool (Fig. 3m, n). Collectively, these data suggest that the divergent (O-Δ) tRBD-NTD not only recruits new Omicron-specific B cells but also expands broadly reactive populations (Delta+ Omicron+; Delta+CoV-1+; multi-binders), providing a single-cell mechanistic basis for the observed increase in functional breadth (Fig. 3g-h). Leveraging the unique design of the O-Δ tRBD-NTD, we show that breadth arises not from accumulating variant-specific responses but from preferentially expanding rare cross reactive B cell clones. This establishes a new, experimentally validated route to engineer broader sarbecovirus immunity.

### Single-cell BCR sequencing shows the divergent (O-Δ) tRBD-NTD recruits clonally distinct, low-SHM cross-reactive B cells

We sought to determine whether the nature of clonal selection of multi-binding B cells differs between matched (Δ-Δ) and divergent (O-Δ) tRBD-NTD vaccines. Specifically do multi-binding B cells arise by the same paths (i.e. via somatic hypermutation or direct recruitment of germline-encoded B cells) in both formats, or does antigen display change the selection dynamics? To answer this, we profiled the clonal distribution by single-cell BCR sequencing of antigen-specific B cells from draining LNs (inguinal and iliac) of immunised mice. Cohorts (n = 12 per vaccine) received two doses of either matched (Δ-Δ) tRBD-NTD or divergent (O-Δ) tRBD-NTD vaccine. Lymph nodes were processed individually, indexed with sample tags and then pooled for cell sorting. Antigen-specific B cells were sorted by multicolour baits into three tiers of polyreactivity: (I) single-binding (Delta only), (ii) mid cross-reactive (Delta+ Wuhan+ or Delta+CoV-1+), and (iii) multi-binding (Delta+Wuhan+CoV-1+) (Fig. 5a). To ensure that cross-reactivity metrics were not artificially driven by inclusion of Omicron in the divergent (O-Δ) tRBD-NTD vaccine, Omicron probes were excluded from the sequencing sorts. Sorted cells were stained with oligo-tagged surrogate antibodies (CD19 and B220) to retain tier identity (Figure 5a; surrogate staining confirmed in Supplementary Fig. 1a) before loading onto BD Rhapsody single-cell separation cartridge. Whole transcriptomic single-cell sequencing, V(D)J sequencing, clone calling and somatic hypermutation quantification were performed on a single cell level.

**Figure 5.**
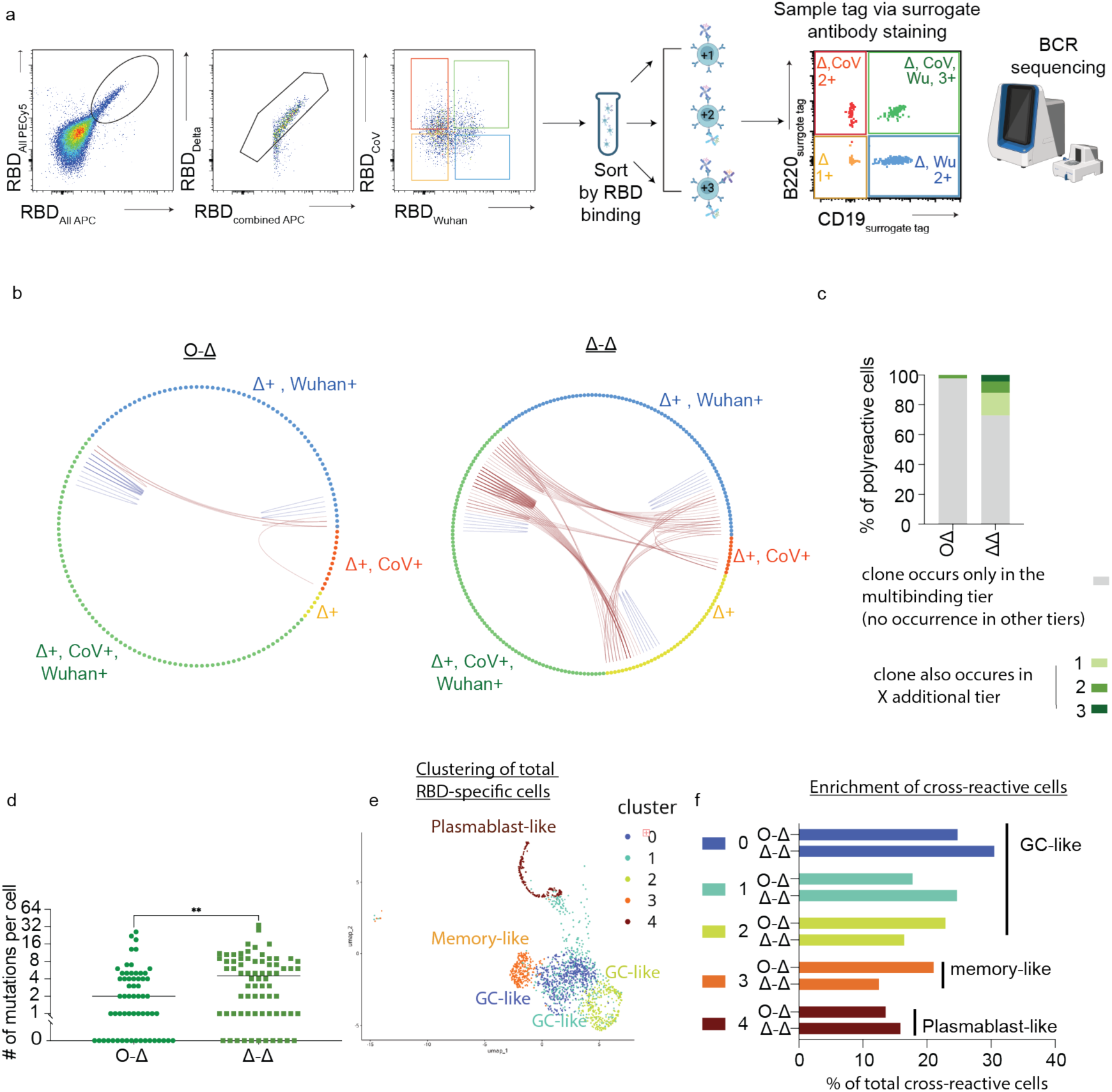
Divergent (O-Δ) tRBD-NTD promotes clonal separation and lower SHM among broadly reactive B cells. Mice were immunised twice, 3 weeks apart and draining lymph nodes were harvested 7 days after the second shot. **a**, Schematic of sorting and sequencing strategy. Antigen-specific B cells were isolated by flow cytometry using multicolour RBD baits, each tier was indexed with oligo-tagged surrogate anti-B220/CD19 antibodies, pools were combined, and single-cell BCR sequencing performed. **b**, Clonal network plots for divergent (O-Δ) and matched (Δ-Δ) tRBD-NTD immunised mice. Each node represents one B cell (colour = binding tier: Yellow is single binding Δ+; red is mid cross-reactive Δ+CoV-1+; blue is mid cross-reactive Δ+Wuhan+; green is multibinding Δ+CoV-1+Wuhan+). Clonal network plots show pooled cells from 12 mice in each group. Lines connect clonally related cells (same V(D)J clone) within an animal. Blue lines: within-tier relationships; red lines: across-tier relationships. **c,** Clone occurrence across tiers for multibinding clones. Bars show the fraction occurring in 0, 1, 2, or 3 additional tiers (0= unique to multibinders; 3 = present in all other tiers) after matched (Δ-Δ) vs divergent (O-Δ) tRBD-NTD immunisation. **d,** Somatic hypermutation per B cell among multi-binders in Δ-Δ vs O-Δ cohorts. **e,** UMAP of antigen-specific B cells from all treatment groups and all mice. **f,** B cell state enrichment plot of multi-binding B cells from matched (Δ-Δ) and divergent (O-Δ) cohorts. Sequencing was performed on pooled cells (n = 12 mice per group), clonal relationships were analysed within each mouse separately. Statistical analysis using a Mann-Whitney test. P value ** is <0.01

We addressed clonal relatedness within each individual animal across the three binding tiers. Specifically, we assessed whether shared clones were found between B cells in different tiers of polyreactivity. In the Network plots (Fig. 5b), nodes represent single B cells (coloured by tier) and lines indicate clonal relationships. Blue lines connect clonally related B cells within the same tier of polyreactivity; red lines connect B cell clones across polyreactive tiers (e.g., a single binding cell clonally related to a multi-binding B cell). In divergent (O-Δ) tRBD-NTD recipients, multi-binding B cell clones were largely clonally distinct from B cells in single-binding and mid cross-reactive tiers of polyreactivity, with minimal cross-tier overlap (only ∼2.3% of multi-binders shared with other tiers of polyreactivity) (Fig. 5b). In contrast, in matched (Δ-Δ) tRBD-NTD recipients, a substantial fraction of multi-binding clones (∼27.2 %) also appeared in single-binding or mid cross-reactive tiers of polyreactivity (Fig. 5b, c), indicating lineage sharing. This pattern is quantified in Fig. 5c, the Δ-Δ cohort shows a greater fraction of multi-binding clones occurring in one to three additional tiers (1-3), whereas the O-Δ cohort is enriched for clones occurring in no additional tiers (0). These results are consistent with breadth arising within shared lineages via maturation in Δ-Δ, and breadth arising by selection of distinct cross-reactive clones in O-Δ.

We next quantified somatic hypermutation (SHM) across binding tiers. SHM frequencies were similar between vaccines for single-binding and mid cross-reactive tiers of polyreactivity (Supplementary Fig 1b). In contrast, a significant difference emerged among multi-binding B cells: multi-binders from the matched (Δ-Δ) tRBD-NTD immunised mice contained more highly mutated sequences, whereas multi-binders from the divergent (O-Δ) tRBD-NTD immunised mice were enriched for low-SHM sequences (Fig. 5d). These patterns indicate that the divergent (O-Δ) tRBD-NTD vaccine directly expands cross-reactive clones with low mutational burden, as seen in the multi-binding cells. This is consistent with preferential recruitment of B cells biased towards cross-variant specificities in the divergent tRBD-NTD, whereas the matched (Δ-Δ) tRBD-NTD more often yields cross-reactivity after iterative mutation/selection. Together, these data suggest that divergent vaccine supports the generation of breadth by selection of clonally unique precursors, providing a more efficient pathway to breadth.

We sought to address whether multi-binding B cells from the two vaccine formats occupy different B cell states. Whole transcriptome analysis revealed four major clusters (Fig. 5e). Clusters 0-2 exhibited GC-like features (*Bcl6, Cd95, Aicda* high); Cluster 3 expressed *Cd38,* consistent with memory-like B cells; and Cluster 4 expressed high levels of *J-chain* and *Xpb1*, consistent with plasmablasts (Supplementary figure 1c). In line with SHM findings, divergent (O-Δ) tRBD-NTD multi-binders were less represented in GC-like clusters, and more in memory-like cluster (Fig. 5f, Supplementary figure 1d), whereas matched Δ-Δ multi-binders were enriched in GC-like clusters (Fig. 5f). These data support a model in which multi-binding cells generated by the divergent vaccine are phenotypically distinct, and earlier in the GC selection trajectory than those arising after the matched (Δ-Δ) tRBD-NTD vaccine. Collectively, these data reveal that antigen geometry can rewire clonal selection. The divergent (O-Δ) immunogen produces breadth primarily by selecting clonally unique, low-SHM, germline-competent B cells that recognise conserved determinants, representing a rapid path to cross-reactivity. By contrast, a matched tRBD-NTD that duplicates every epitope tends to produce breadth via extended GC-mediated maturation of variant-specific precursors. This mechanistic distinction is both conceptually and practically important, because it shows that vaccine design can be used not only to increase breadth, but to control the path by which breadth is achieved, a fundamentally different strategy from existing multivalent or mosaic approaches.

## Discussion

Here we describe a structurally distinctive, compact mRNA immunogen in which two antigenically divergent RBDs are cis-linked on a single membrane-anchored scaffold, producing nanoscale proximity of duplicated conserved epitopes. This single-ORF tRBD-NTD design simplifies manufacture relative to multi-protein or nanoparticle platforms and preserves an NTD module to enhance CD4⁺ T-cell help, following the basic design principles established in Montgomerie et al. Crucially, going beyond the previous design, this tandem-RBD design increases the local copy number of conserved surfaces and is likely to enable two avidity gains at the BCR: intramolecular bivalency, and intermolecular BCR cross-linking^38,39^ across repeated conserved patches. Both mechanisms are well-supported by work showing that nanoscale antigen organisation, valency, and spacing govern BCR clustering, signalling and subsequent T_FH_-mediated selection in GCs^40-42^.

In this study, our key mechanistic finding was that epitope density can fundamentally alter clonal selection. Using single-cell tetramer profiling, paired V(D)J sequencing and transcriptome analysis, we show that broadly reactive anti-RBD clones elicited by the divergent construct exhibited lower SHM than those from the matched (Δ-Δ) tRBD-NTD vaccine. This finding fits with reports that some bnAbs can be germline-encoded, including RBD- and NTD-directed specificities^43-45^, while others require SHM to achieve breadth^5,6,46^. Importantly, unlike HIV bnAbs, SARS-CoV-2 breadth does not generally demand extreme or unusual mutation; both low-SHM and highly mutated broad specificities arise after standard immunisation or infection. The central challenge is therefore preferentially recruiting and expanding the right clones. Two independent single-cell readouts support a breadth-by-selection mechanism for our design.

First, multicolour bait/flow cytometry revealed expansion of variant-specific (e.g., Omicron-only) cells and of co-binders that recognise both vaccine RBDs. Critically, we also observed Delta+ Wuhan and Delta+CoV-1+ co-binding despite that these strains were absent from the vaccine itself, which is a pattern consistent with recruitment of clones targeting shared, functionally constrained RBD surfaces^5,9^. Second, single-cell BCR-seq showed that multi-binding cells in the divergent arm were clonally distinct from variant-specific pools and low-SHM, whereas in the matched arm, multi-binders more often shared lineage with single-binders and carried higher SHM, consistent with breadth acquired via maturation^5,6,46^. Together, these data support a model in which duplicating conserved patches by placing two divergent RBDs in intramolecular, proximity biases GC competition toward germline-biased B cells that recognise those patches, rather than relying on prolonged affinity maturation. Our approach presents an alternative to classical germline-targeting (precursor-targeting) strategies^38,39,47-49^. Instead of epitope-engineered scaffolds designed to bind germline precursors, we specifically raised the local copy number of naturally conserved RBD surfaces lowering the activation threshold for germline-encoded, conserved-epitope specificities to win in the GC^38^.

Rapid antigenic diversification of SARS-CoV-2 has repeatedly narrowed the coverage of first-generation vaccines and monoclonals, underscoring that durable protection will require responses centred on epitopes with constrained mutational tolerance. Several reports have shown that bnAbs to conserved RBD surfaces, such as overlapping ACE2-contact (“receptor-mimicry”) patches or conserved non-RBM “silent-face/site-V” footprints do exist but are uncommon within typical vaccine or infection repertoires, highlighting a selection bottleneck rather than biological impossibility^12,15,16^. Strategies to enhance breadth span phylogeny-guided or mosaic nanoparticle displays, consensus/computational antigens, chimeric/multivalent spikes (including S2-focused), and epitope-focused immunogens. Consensus and designed antigens can broaden serum cross-neutralisation but still face immunodominance and manufacturing complexity as sequences evolve^12,22-25^. Mosaic and phylogeny-guided RBD nanoparticles boost cross-reactive titres by presenting diverse sequences at high valency, yet require co-production and controlled stoichiometry of multiple proteins^14-18^. Chimeric or S2-multivalent spikes can expand breadth but may channel responses toward immunodominant, variably protective specificities^14,18,22-25^. Epitope-focused scaffolds precisely target conserved sites but often trade off precision with poor manufacturability and expression, and breadth can be fragile if the targeted surfaces are not sufficiently constrained^19-21^. In contrast, our single-molecule, membrane-anchored tRBD-NTD minimises payload complexity while increasing conserved-epitope density by design, which we envisage as an approach orthogonal to (and potentially combinable with) these platforms. This work offers a manufacturable design rule for next-gen boosters and pan-sarbecovirus candidates. Increasing the copy number of critical conserved epitopes without repeating epitopes that are easily mutated could further increase the fraction of antibodies with bnAb-like specificities, helping move the field from breadth as an emergent average to breadth as an engineered outcome.

While our data demonstrate enhanced breadth of the humoral response in murine models; further studies are needed to evaluate durability, protective efficacy against live virus challenge, and translational relevance in humans. Epitope mapping of cross-reactive clones and structural characterisation of antibodies elicited by divergent tRBD-NTD vaccination would clarify mechanisms of breadth.

Related work using a SARS-CoV-1 and SARS-CoV-2 RBD heterodimer reported improved serum responses across variants^50^, underscoring that co-presentation of divergent RBDs can broaden activity at the population level. Our data extend this concept with single-cell resolution. This complements prior serum-level observations^50^ and provides a mechanism for selecting pairings that bias breadth by selection rather than relying on extensive maturation.

Studies examining serum responses to bivalent mRNA boosters (e.g., BA.4/5) show modest and transient gains against Omicron^10^, and studies of Omicron breakthrough after vaccination reveal different outcomes, in some settings, expansion of cross-binding B cells^51^, in others, Omicron-specific responses that evolve largely independently of ancestral specificities^52^. Together, these findings argue that simply co-delivering variant spikes is insufficient to reliably broaden or prolong immunity without a mechanism that enriches cross-reactive B cells.

Together, these data argue that breadth can be designed into the earliest steps of clonal selection. By cis-linking divergent RBDs, tRBD-NTD concentrates conserved epitopes and preferentially recruits germline-biased B cells that recognise those constraints. This shifts breadth from a stochastic, maturation-dependent outcome to a selectable property of antigen architecture. Implemented in a single-ORF, mRNA format, this provides a simple, scalable design principle for enriching rare, conserved-epitope specificities, that support broad protection against current and future variants.

## Methods

### Vaccine preparation

#### mRNA synthesis

mRNA was synthesised as previously described^53^. Inserts were cloned 5’*Age*I and 3’*EcoR*I restriction sites into a modified pVAX1 vector (Thermo Fisher) containing a T7 RNA polymerase promoter, UTRs and *Not*I restriction site for template linearization. Linearized template was purified using a PCR & DNA Clean up kit (NEB), and *in vitro* transcription was carried out using a HiScribe T7 High Yield RNA Synthesis Kit (NEB). DNA template was removed with DNase I (NEB) prior to precipitation with 6M LiCl at −20 °C for 30 min, followed by centrifugation 13,000g for 30 min at 4 °C. A 5’ cap and 3’ polyA tail were added to the RNA using the Vaccinia Capping System and *Escherichia coli* polyA polymerase, respectively (NEB). RNA integrity and polyA tail were confirmed using agarose gel electrophoresis and RNA ScreenTape analysis (Agilent 4200 TapeStation, Agilent Technologies Ltd.).

#### Encapsulation

##### mRNA-LNP Formulation

mRNA was encapsulated as previously described^53^. Lipids composed of a mixture of ALC-0315, DSPC, cholesterol and ALC-0159 in a molar ratio of 46.3/9.4/38.5/1.6, respectively, were dissolved in ethanol at 10 mg/mL. mRNA was dissolved in 100 mM pH 5.2 acetate buffer at 140 µg/mL. The organic and aqueous phases were mixed on a NanoAssemblr Ignite™ using a NX Gen™ cartridge using a total flow rate of 12 mL/min and flow rate ratio of 3:1 to achieve a final N/P ratio of 6. The resulting suspension was diluted 15-fold in PBS buffer and concentrated using an Amicon centrifugal concentrator with MWCO 100kDa. mRNA-iLNP suspensions were stored at 4 °C until use.

##### mRNA-LNP Characterisation

Particle size, polydispersity index (PDI) and zetapotential of mRNA-LNPs diluted 10-fold in PBS were analysed on a Zetasizer Nano ZS (Malvern, UK) at 25 °C. mRNA encapsulation efficiency and concentration were determined using the Quant-it™ RiboGreen RNA Assay. Encapsulation efficiency was calculated in duplicate as (F_T_ - F_0_)/F_T_ were F_T_ and F_0_ are the amount of mRNA quantified in presence and absence of 1 % triton X-100, respectively.

##### Typical Results

| Lipids | N/P | Size<sup>a</sup><br>nm | PDI<sup>a</sup> | Encapsulation<br>efficiency<sup>b</sup> % | Zeta-potential<sup>a</sup><br>mV |
| --- | --- | --- | --- | --- | --- |
| 46.3 ALC-0315/9.4 DSPC/42.7 cholesterol/1.6 ALC-0159 | 6 | 85 | 0.070 | 85 | 2.8 |

<sup>a</sup>Average of three runs of the same sample. <sup>b</sup>Average of duplicate analysis of the same sample.
<sup>c</sup>Construct length before tailing. <sup>d</sup>Determined by RiboGreen assay.

#### tRBD-NTD structure

The structure of the tRBD-NTD based on our previously described vaccine^30^. Briefly RBDs were linked by a single glycine linker, a glycine and a serine were inserted between the C-terminus RBD and the NTD, which was translated to the C-terminus. The TMD is WYIWLGFIAGLIAIVMVTIMLCCMTSCCSCLKGCCSCGSCCKFD; an arginine and glycine link the NTD and the TMD.

### Immunizations

Specific pathogen-free C57BL/6J mice were bred and housed at the Malaghan Institute of Medical Research. Breeding pairs originated from The Jackson Laboratory. Sex- and age-matched mice (9– 14 weeks old, matched within 2 weeks) were used in all experiments. Procedures were approved by the Victoria University of Wellington Animal Ethics Committee. Mice were anaesthetised by intraperitoneal injection of ketamine/xylazine (ProVet NZ) and immunised intramuscularly in both hind legs (50 µL per leg) with 5 µg mRNA–LNP vaccine diluted in DPBS (Gibco) using 0.5 mL insulin syringes (BD Biosciences). Boosts were given 21 days after the primary dose, and the response was measured one-week post-boost.

### tRBD-NTD modelling

The potential for a single IgG antibody molecule to interact with out tandem RBD immunogens was assessed using molecular modelling. First, AlphaFold3^54^was used to predict structures for the matched tRBD-NTD (Δ-Δ) and the divergent tRBD-NTD (O-Δ). Next, we used AlphaFold3 to assess interactions between the O-Δ tRBD-NTD immunogen and various monoclonal antibodies. The experimentally determined structures of the broadly neutralising anti-RBD antibodies ZCP3B4 (PDB 8K19), VacBB-551 (PDB 8GS9), BD55-1205 (PDB 8XE9), and K4-66 (PDB 9II9) contained individual Fab domains but not full-length IgG molecules. Therefore, we took the variable regions of these antibodies and introduced them into the sequence of a full-length IgG1 antibody that had an experimentally determined structure (PDB 1IGY). In each of the four cases, the chimeric, full-length IgG sequence was uploaded to the AlphaFold3 web server along with the sequence of the divergent O-Δ tRBD-NTD immunogen. The input to AlphaFold3 therefore comprised five polypeptide chains (two antibody light chains, two antibody heavy chains, and one RBD vaccine chain). The resulting predictions from AlphaFold3 showed clear interactions between each immunogen and a correctly-assembled IgG molecule. As a control, we uploaded the sequence of the immunogen together with the unmodified sequence from PDB entry 1IGY. This experimentally determined structure is of a full-length monoclonal antibody that recognises phenobarbital^35^ AlphaFold3 predicted no interaction between this “irrelevant antibody” and our immunogen.

The modelled structures were visualised with UCF ChimeraX version 1.9^55^and the Predicted Aligned Error (PAE) scores from AlphaFold3 are presented as PAE plots. ISOLDE version 1.10.1^56^ was used to reposition the antibody Fc domain above the Fab domains and the NTD below the RBDs for visual clarity. During the ISOLDE simulations, the Fab and RBD domains were pinned to prevent disruption of any RBD-antibody interfaces; at no point were these domains altered. On the other hand, the PAE plots of the vaccine NTD domain did not show interactions with any other domain, and the antibody Fc region of each heavy chain interacted only with its corresponding Fc heavy chain counterpart. Therefore, we used ISOLDE to rotate the Fc domain around the antibody hinge region between the Fc and Fab domains, and to rotate the NTD around the flexible glycine linker between the second RBD domain and the NTD domain. The simulations were informed by PAE plots to ensure that all predicted antibody-immunogen interactions were maintained.

### Antibody assessment

#### Conventional ELISA

RBD-specific serum antibodies were measured by ELISA as previously described^57^. Briefly, Nunc MaxiSorp™ 96-well plates (ThermoFisher Scientific) were coated overnight at 4 °C with yeast-expressed RBD protein. Plates were washed with DPBS + 0.05% Tween-20 (Sigma-Aldrich) and blocked with 10% FBS (Gibco) in DPBS for 1h at room temperature (RT). Serially diluted serum samples were added and incubated for 2h, followed by HRP-conjugated goat anti-mouse IgG (Invitrogen) for 1h at RT. After washing, TMB substrate (OptEIA, BD Biosciences) was applied, reactions were stopped with 2M H₂SO₄, and absorbance was measured at 450 nm (Perkin Elmer EnSpire 2300). End-point titres were determined using the t-statistic method^58^ in GraphPad Prism.

#### Binding affinity assay

Antibody affinity was evaluated using a urea dissociation ELISA. The standard ELISA protocol was followed with an additional step to remove low-affinity antibodies. Diluted serum samples were plated in duplicate for standard and urea-treated conditions. Following the 2h incubation, plates were washed and treated with either 6M urea (Sigma-Aldrich) or DPBS for 30 min at 37 °C to dissociate weakly bound antibodies. Plates were washed and developed as described above. The affinity index was calculated as the ratio of OD450 values between urea-treated and standard wells, measured at the dilution corresponding to 80% of the maximal signal in untreated samples, ensuring analysis within the linear portion of the response curve.

#### Pseudovirus neutralisation assay

Neutralising activity was quantified using a lentiviral-based SARS-CoV-2 pseudovirus assay adapted from Crawford et al. HEK293T-hACE2 cells (2.5 × 10⁴/well) were seeded in poly-D-lysine–coated, 96-well white-walled plates and incubated at 37°C, 5% CO₂ for 24 h. Mouse sera were heat-inactivated (56 °C, 30 min) and serially diluted (1:10, 1:5) in DMEM supplemented with puromycin (2 µg/mL) and geneticin (500 µg/mL). Equal volumes of pseudotyped viral particles were added and incubated for 1h at 37 °C before addition of polybrene (5 µg/mL). After 72h, viral entry was quantified using the Steady-Luc Firefly HTS assay (Promega). Supernatants were removed, luciferin reagent (1:1 with DMEM) was added, and plates were incubated for 5min with shaking. Luminescence was read on a Perkin Elmer EnSpire 2300. Neutralisation titres (NT₅₀) were calculated using non-linear regression in GraphPad Prism.

### Flow cytometry

#### Confirming ACE2 binding

Expi293 cells were transfected with plasmids encoding matched or divergent (O-Δ) tRBD-NTDs, or the full-length SARS-CoV-2 spike using Expifectamine according to the manufacturer’s instructions. Protein expression was assessed 4 days post-transfection. Non-specific staining was blocked with 20% FCS in DPBS for 1 hour. Cells there then stained with hACE-Fc (made in house), Mouse anti-Human IgG1 Fc Secondary Antibody, Alexa Fluor™ 488 (ThermoFischer Cat # A-10631).

#### Tetramer formation

The BirA tag was added to the N-terminus of the RBDs, connected by a 6xHIS tag. SARS-CoV-2 Wuhan, Delta, and CoV-1 RBD construct was codon optimized for human cell expression, synthesized and sequence-verified by Gene Universal (Geneuniversal.com). Both tags are cleavable by HRV-3C protease (LEVLFQ/GP) (made in house) during the purification process. HEK293 cells lacking N-acetylglucosaminyltransferase I (GnTI-) activity (HEK293 GnTI-) were obtained from American Type Culture Collection (ATCC, Manassas, VA). HEK293 GnTI-cells were cultured in DMEM supplemented with 5% FBS at 37°C and 5% CO_2_. Stable cell lines were made for all constructs by selecting for resistance to Geneticin (G418) (Sigma) using DMEM supplemented with 5% FBS and 500 µg/ml of G418. Resistant clones were isolated using Pyrex cloning rings (Merck) and stable expression of the proteins was tested by Western blot using anti-FLAG antibody (M2) (Sigma). Clones expressing high levels of the protein were amplified, frozen, and used in large-scale protein production. Milligram-scale protein production was performed in triple layer cell culture flasks. Secreted proteins were affinity purified from cell culture media using Protein A-CaptivATM PriMAB (RepliGen, Waltham, MA) in 150 mM NaCl, 50 mM Tris, pH 8.0 or anti-FLAG® M2 affinity resin (Sigma), and subsequently cleaved with HRV 3C protease (made in house) to remove the FLAG and Fc fragments. Residual HRV-3C protease and Fc fragments were removed from the protein of interest by a coupled negative purification step using GSTrap™ HP and HiTrap MabSelect PrismA (Cytiva) columns, respectively, in PBS. Purified protein was stored at 4°C and used within a week or aliquoted and flash frozen in liquid nitrogen and stored at -80°C.

RBDs were biotinylated using the Sigma Aldrich Enzymatic Protein Biotinylation kit (CS0008). Biotinylation was confirmed by ELISA. Commercially available NP-OVA-biotin (Creative Diagnostics) was used as an irrelevant bait negative control. Separate reactions were set up to add two fluorescently labelled streptavidin (SAV) to RBD and NP-OVA. SAV was added at a 1:20 protein to SAV ratio. One fifth of the fluorescently labelled SAV was added to the protein at a time, with 20-minute incubations between applications, on ice. When all the SAV was added, 4 mM free biotin (Sigma) in PBS was added at a 1:1 volume ratio, to saturate free SAV, for 30 minutes. The tetramers were then spun at max. speed at 4°C for 10 minutes. NP-OVA formed tetramers with SAV-PE, critically providing an opportunity to remove PE specific B cells. All RBDs formed tetramers with SAV PE-Cy5 and SAV APC-Cy7. RBDs then formed tetramers with unique SAVS; Delta PECy5.5, Omicron BV605; CoV PeCy7, Wuhan PECS595. Cells were stained with 1:50 dilution per tetramer in 4mM free biotin for 30 minutes. Specific cells were identified as NP-OVA negative. All samples were collected on a Cytek® Aurora spectral flow cytometer (Cytek®). Compensation was performed in each experiment using UltraComp eBeads (Invitrogen). Analysis was performed using FlowJo version 10 (BD biosciences).

#### B cell surface staining

##### Preparation of tissues for immunological assessment

C57BL/6J mice were euthanised, and blood, spleens, and inguinal and iliac lymph nodes were collected. Blood was obtained by cardiac puncture into S-Microvette® serum gel tubes (Sarstedt), centrifuged at 10,000 × g for 5 min, and serum aliquots were stored at −80 °C. Spleens were processed into single-cell suspensions by gently teasing through 70 µm strainers (Falcon) and washing with Iscove’s Modified Dulbecco’s Medium (IMDM; Gibco). Cells were centrifuged at 250 × g for 10 min, red blood cells lysed (Qiagen), washed at 300 × g for 4 min, resuspended in RPMI-1640 (Gibco) containing 10% heat-inactivated FBS, and filtered through 70 µm strainers. Lymph nodes were similarly dissociated through 70 µm strainers, washed in IMDM, pelleted at 250 × g for 10 min, and resuspended in R10 media. Cell counts were determined using SPHERO AccuCount Blank Particles (Spherotech).

##### Surface staining

Cells were blocked with anti-mouse CD16/32 (clone 2.4G2) for 5 min at room temperature, then stained with Zombie NIR viability dye (BioLegend). Surface staining was performed at 4 °C for 15 min using fluorescent antibody cocktails in PBS with 0.01% sodium azide, 2 mM EDTA, and 2% FBS. Cells were then stained on ice for 30 min with RBD–streptavidin tetramers, fixed, and permeabilised using the eBioscience™ FoxP3/transcription factor buffer set (Invitrogen) before intracellular staining for 1 h. Data were acquired on a Cytek Aurora (SpectroFlo v3.3.0) and analysed using FlowJo v10.10.

### FACs sorting

Cells were stained with fluorescently labelled antibodies against TCRβ, NK1.1, CD11c, B220, IgD, and CD138, along with fluorescent RBD tetramers representing SARS-CoV-2 Delta, ancestral (Wuhan-Hu-1), and SARS-CoV-1 variants. Delta RBD-binding B cells were identified as B220⁺NK1.1⁻CD11c⁻TCRβ⁻IgD⁻CD138⁻ cells that bound both pooled RBD tetramer fluorophores and were positive for Delta RBD. Within this Delta⁺ population, cells were further subdivided based on cross-reactivity. B cells binding only the Delta RBD were classified as into three tiers of polyreactivity: (I) single-binding (Delta only), (ii) mid cross-reactive (Delta+ Wuhan+ or Delta+CoV-1+), and (iii) multi-binding (Delta+Wuhan+CoV-1+). Each subset was subsequently labelled with a unique combination of surrogate oligo-tagged TotalSeq antibodies (BioLegend) to enable post hoc deconvolution following single-cell sequencing.

### RNAseq

cDNA Libraries were prepared using the Mouse TCR/BCR Full Length, mRNA Whole Transcriptome Analysis (WTA), BD® AbSeq, and Sample Tag Protocol (BD Biosciences, 23-24367(01)), mRNA WTA and Sample Tag Library kits according to the manufacturer’s instructions. The quality of the final libraries was assessed using a TapeStation 4150 (Agilent) with High Sensitivity D5000ScreenTape and quantified using a Quantus Fluorometer (Promega). BCR, WTA, AbSeq and Sample Tag libraries were diluted to 2 nM and mixed at a ratio of 52.2%, 42.0%, 4.2%, and 1.3% respectively. The libraries were loaded on an P1 flow cell (2 × 150 cycle) and paired-end sequenced at >50,000 reads per cell depth on a Novaseq 1000 Sequencer (Illumina) at Ramaciotti Centre for Genomics, UNSW, Sydney, Australia. PhiX was added to the sequencing run to compensate for the low complexity library. (Additional reads were obtained for the AbSeq and Sample Tag library to make up for the imbalance of read depth among Sample Tags and AbSeq antibodies).

### Analysis

Raw single-cell reads from tagged mice were processed through three parallel pipelines to extract complementary information: 1) gene expression - reads were aligned to the mouse Gencode M36 transcriptome using bowtie2 to generate per-gene transcript counts; 2) sample/Abseq tagging - reads were processed with the Velsera / BD Rhapsody Sequence Analysis Pipeline to quantify Sample Tag indices (vaccine group) and AbSeq antibody-derived tag counts (surrogate labelling of B cell polyreactivity tiers); and 3) BCR sequences - reads were analysed with TRUST4 to reconstruct BCR sequences and assign BCR clonotypes. These intermediate datasets were merged using BD cell barcode sequences as a primary key, resulting in a single combined dataset that categorised each cell based on its vaccine type (via AbSeq tags) and associated BCR clonotype. Circos plots were created that linked cells to other cells that shared the same clonotype. We removed genes with < 250 total molecules across the dataset and cells with <250 total molecules, as well as cells with >= 35% mitochondrial or >= 90% immunoglobulin reads. Expression values were normalised and scaled with SCTv2 using v5.3.0 and cells were clustered at a resolution of 0.8^59^. Four of the nine identified clusters had negligible BCR transcription and were excluded.

### Statistics Quantification and Statistical Analysis

Statistical analyses were performed using GraphPad Prism v.10.4.1 (532) on pooled data from 2-4 independent experiments each with 5 mice per group. Statistical comparing only two groups were performed using Mann Whitney test. Statistical difference between more than 2 groups was determined using a one-way ANOVA, with no matching, followed by a Fishers’s post-test. The statistical difference between more than 1 variant for more than 2 groups was determined using a two-way ANOVA with a Fishers post-test. P values of <0.05 were considered statistically significant. Figures were generated in Adobe Illustrator and BioRender.com.

### Key Resources

**Table 1:** Antibodies.

| Item | Source | Identifier (catalog #) |
| --- | --- | --- |
| **Fluorescent reagents** |  |  |
| B220 BUV737 (clone RA3-6B2) | BD Biosciences | 612838 |
| CD11c BV786 (clone HL3) | BD Biosciences | 563735 |
| CD45 BUV395 (clone 30-F11) | Invitrogen | 363-0451-82 |
| CD45 BV711 (clone 30-F11) | BioLegend | 103147 |
| IgD APC H7 (clone 11-26c.2a) | BD Biosciences | 565348 |
| NK1.1 BUV805 (clone PK136) | BD Biosciences | 741926 |
| SAV APC | BD Biosciences | 554067 |
| SAV APC Cy7 | BioLegend | 405208 |
| SAV BV605 | BioLegend | 405229 |
| SAV PE | BioLegend | 405204 |
| SAV PE CF594 | BD Biosciences | 562284 |
| SAV PE Cy5 | BioLegend | 405205 |
| SAV PE Cy5.5 | Invitrogen | SA1018 |
| SAV PE Cy7 | BD Biosciences | 557598 |
| **ELISA reagents** |  |  |
| anti-HIS tag HRP | BioLegend | J099B12 |
| Goat anti-mouse total IgG HRP | Invitrogen | G21040 |
| Monoclonal mouse anti-spike IgG1 clone 43 antibody | SinoBiological | 40591-MM43 |
| Mouse anti-human IgG (Fc) HRP | BioRad | MCA647P |

**Table 2:** Labware.

| Item | Source | Identifier (catalog #) |
| --- | --- | --- |
| 1mL syringe | BD Biosciences | 302113 |
| 0.5mL insulin syringe | BD Biosciences | 326105 |
| 70µM cell strainer | Falcon® | BDAA352350 |
| 96-well white polystyrene microplates | Corning® | CLS3610 |
| Microvette serum gel tubes | Sarstedt | SARS20.1344 |
| Nunc MaxiSorp™ 96-well ELISA microplates | ThermoFisher Scientific | 442404 |
| S-Microvette® serum gel tubes | Sarstedt | 20.1344 |

**Table 3:** Reagents.

| Item | Source | Identifier catalog # |
| --- | --- | --- |
| Dulbecco's phosphate-buffered saline (DPBS), no calcium, no magnesium | Gibco™ | 14190250 |
| DMEM (Dulbecco's Modified Eagle Medium), high glucose, GlutaMAX™ supplement, pyruvate | Gibco™ | 10569044 |
| DPBS powder, no calcium, no magnesium | Gibco™ | 21600069 |
| eBioscience™ FoxP3/transcription factor fixation/permeabilisation kit | Invitrogen | 00-5521-00 |
| Enzymatic Protein Biotinylation Kit | Sigma-Aldrich | CS0008 |
| Fc Block CD16/32 (clone 2.4G2) | Made in-house | - |
| Fetal bovine serum (FBS) | Gibco™ | 10091-148 |
| Geneticin™ | Gibco™ | 10131035 |
| hACE-Fc | Made in-house | - |
| HEK293 GnTI | American Type Culture Collection ATCC, Manassas, VA | CRL-3022 |
| Iscove's Modified Dulbecco's Medium (IMDM), GlutaMAX™ supplement | Gibco™ | 31980097 |
| OptEIA™ TMB substrate reagent set | BD Biosciences | 555214 |
| Poly-D-lysine | Gibco™ | A3890401 |
| Puromycin dihydrochloride | Gibco™ | A1113802 |
| Red blood cell lysis solution | Qiagen | 158904 |
| Roswell Park Memorial Institute (RPMI) 1640 medium | Gibco™ | 11875-119 |
| Steady-Luc Firefly HTS Assay Kit | Biotium | BIT30028-3 |
| Tween® 20 | Sigma-Aldrich | P9416 |
| Urea powder | Sigma-Aldrich | U5378-500G |

## Supporting information

Plain text sequence

Pain text sequence

