## Supplementary material for "Engineering antigenic breadth against SARS-CoV-2 by pairing divergent RBDs within a single mRNA immunogen": Plain text sequence

gactcttcgcgatgtacgggccagatatacgcgttgacattgattattgactagttattaatagtaat
caattacggggt
cattagttcatagcccatatatggagttccgcgttacataacttacggtaaatggcccgcctggctga
ccgcccaacgac
ccccgcccattgacgtcaataatgacgtatgttcccatagtaacgccaatagggactttccattgacg
tcaatgggtgga
ctatttacggtaaactgcccacttggcagtacatcaagtgtatcatatgccaagtacgccccctattg
acgtcaatgacg
gtaaatggcccgcctggcattatgcccagtacatgaccttatgggactttcctacttggcagtacatc
tacgtattagtc
atcgctattaccatggtgatgcggttttggcagtacatcaatgggcgtggatagcggtttgactcacg
gggatttccaag
tctccaccccattgacgtcaatgggagtttgttttggcaccaaaatcaacgggactttccaaaatgtc
gtaacaactccg
ccccattgacgcaaatgggcggtaggcgtgtacggtgggaggtctatataagcagagctctctggcta
actagagaaccc
actgcttactggcttatcgaaattaatacgactcactatagggagacccaagctggCTAGCcgcaggc
cccgcccgggac
tcccctgcggtccaggccgcgccccgggctccgcgccagccaatgagcgccgcccggccgggcgtgcc
cccgcgccccaa
gcataaaccctggcgcgctcgcggcccggcactcttctggtccccacagactcagagagaacccaccA
CCGGTgccaccA
TGTTCGTTTTCCTGGTGCTGCTGCCTCTGGTGTCCAGCACCCGGGTGCAGCCCACCGAATCCATCGTG
CGGTTCCCCAAT
ATCACCAATCTGTGCCCCTTCGGCGAGGTGTTCAATGCCACCAGATTCGCCTCTGTGTACGCCTGGAA
CCGGAAGCGGAT
CAGCAATTGCGTGGCCGACTACTCCGTGCTGTACAACTCCGCCAGCTTCAGCACCTTCAAGTGCTACG
GCGTGTCCCCTA
CCAAGCTGAACGACCTGTGCTTCACAAACGTGTACGCCGACAGCTTCGTGATCCGGGGAGATGAAGTG
CGGCAGATTGCC
CCTGGACAGACAGGCAAGATCGCCGACTACAACTACAAGCTGCCCGACGACTTCACCGGCTGTGTGAT
TGCCTGGAACAG
CAACAACCTGGACTCCAAAGTCGGCGGCAACTACAATTATCGGTACCGCCTGTTCCGGAAGTCCAATC
TGAAGCCCTTCG
AGCGGGACATCTCCACCGAGATCTATCAGGCCGGCAGCAAGCCTTGTAACGGCGTGGAAGGCTTCAAC
TGCTACTTCCCA
CTGCAGTCCTACGGCTTTCAGCCCACAAATGGCGTGGGCTATCAGCCCTACAGAGTGGTGGTGCTGAG
CTTCGAACTGCT
GCATGCCCCTGCCACAGTGTGCGGCCCTAAGAAAAGCACCAATCTCGTGAAGAACAAAGGGCGGGTGC
AGCCCACCGAAT
CCATCGTGCGGTTCCCCAATATCACCAATCTGTGCCCCTTCGGCGAGGTGTTCAATGCCACCAGATTC
GCCTCTGTGTAC
GCCTGGAACCGGAAGCGGATCAGCAATTGCGTGGCCGACTACTCCGTGCTGTACAACTCCGCCAGCTT
CAGCACCTTCAA
GTGCTACGGCGTGTCCCCTACCAAGCTGAACGACCTGTGCTTCACAAACGTGTACGCCGACAGCTTCG
TGATCCGGGGAG
ATGAAGTGCGGCAGATTGCCCCTGGACAGACAGGCAAGATCGCCGACTACAACTACAAGCTGCCCGAC
GACTTCACCGGC
TGTGTGATTGCCTGGAACAGCAACAACCTGGACTCCAAAGTCGGCGGCAACTACAATTATCGGTACCG
CCTGTTCCGGAA
GTCCAATCTGAAGCCCTTCGAGCGGGACATCTCCACCGAGATCTATCAGGCCGGCAGCAAGCCTTGTA
ACGGCGTGGAAG
GCTTCAACTGCTACTTCCCACTGCAGTCCTACGGCTTTCAGCCCACAAATGGCGTGGGCTATCAGCCC
TACAGAGTGGTG

GTGCTGAGCTTCGAACTGCTGCATGCCCCTGCCACAGTGTGCGGCCCTAAGAAAAGCACCAATCTCGT
GAAGAACAAAGG
AAGTAGCCAGTGTGTGAACCTGCGAACCAGAACACAGCTGCCTCCAGCCTACACCAACAGCTTTACCA
GAGGCGTGTACT
ACCCCGACAAGGTGTTTAGATCCAGCGTGCTGCACTCTACCCAGGACCTGTTCCTGCCTTTCTTCAGC
AACGTGACCTGG
TTCCACGCCATCCACGTGTCCGGCACCAATGGCACCAAGAGATTCGACAACCCCGTGCTGCCCTTCAA
CGACGGGGTGTA
CTTTGCCAGCACCGAGAAGTCCAACATCATCAGAGGCTGGATCTTCGGCACCACACTGGACAGCAAGA
CCCAGAGCCTGC
TGATCGTGAACAACGCCACCAACGTGGTCATCAAAGTGTGCGAGTTCCAGTTCTGCAACGACCCCTTC
CTGGATGTCTAC
TACCACAAGAACAACAAGAGCTGGATGGAGTTCGGTGTGTACAGCAGCGCCAACAACTGCACCTTCGA
GTACGTGTCCCA
GCCTTTCCTGATGGACCTGGAAGGCAAGCAGGGCAACTTCAAGAACCTGCGCGAGTTCGTGTTTAAGA
ACATCGACGGCT
ACTTCAAGATCTACAGCAAGCACACCCCTATCAACCTCGTGCGGGATCTGCCTCAGGGCTTCTCTGCT
CTGGAACCCCTG
GTGGATCTGCCCATCGGCATCAACATCACCCGGTTTCAGACACTGCTGGCCCTGCACAGAAGCTACCT
GACACCTGGCGA
TAGCAGCAGCGGATGGACAGCTGGTGCCGCCGCTTACTATGTGGGCTACCTGCAGCCTAGAACCTTCC
TGCTGAAGTACA
ACGAGAACGGCACCATCACCGACGCCGTGGATGCCGGTTGGTACATCTGGCTGGGCTTTATCGCCGGA
CTGATTGCCATC
GTGATGGTCACAATCATGCTGTGTTGCATGACCAGCTGCTGTAGCTGCCTGAAGGGCTGTTGTAGCTG
TGGCAGCTGCTG
CAAGTTCGACTGATGAGAATTCgctggagcctcggtggccatgcttcttgccccttgggcctcccccc
agcccctcctcc
ccttcctgcacccgtacccccgtggtctttgaataaagtctgagtgggcggcagcctgtgtgtgcctg
agttttttccct
cagcaaacgtgccaggcatgggcgtggacagcagctgggacacacatggctagaacctctctgcagct
ggatagggtagg
aaaaggcaggggGCggccgctcgagtctagagggcccgtttaaacccgctgatcagcctcgactgtgc
cttctagttgcc
agccatctgttgtttgcccctcccccgtgccttccttgaccctggaaggtgccactcccactgtcctt
tcctaataaaat
gaggaaattgcatcgcattgtctgagtaggtgtcattctattctggggggtggggtggggcaggacag
caagggggagga
ttgggaagacaatagcaggcatgctggggatgcggtgggctctatggcttctactgggcggttttatg
gacagcaagcga
accggaattgccagctggggcgccctctggtaaggttgggaagccctgcaaagtaaactggatggctt
tctcgccgccaa
ggatctgatggcgcaggggatcaagctctgatcaagagacaggatgaggatcgtttcgcatgattgaa
caagatggattg
cacgcaggttctccggccgcttgggtggagaggctattcggctatgactgggcacaacagacaatcgg
ctgctctgatgc
cgccgtgttccggctgtcagcgcaggggcgcccggttctttttgtcaagaccgacctgtccggtgccc
tgaatgaactgc
aagacgaggcagcgcggctatcgtggctggccacgacgggcgttccttgcgcagctgtgctcgacgtt
gtcactgaagcg
ggaagggactggctgctattgggcgaagtgccggggcaggatctcctgtcatctcaccttgctcctgc
cgagaaagtatc
catcatggctgatgcaatgcggcggctgcatacgcttgatccggctacctgcccattcgaccaccaag
cgaaacatcgca

tcgagcgagcacgtactcggatggaagccggtcttgtcgatcaggatgatctggacgaagagcatcag
gggctcgcgcca
gccgaactgttcgccaggctcaaggcgagcatgcccgacggcgaggatctcgtcgtgacccatggcga
tgcctgcttgcc
gaatatcatggtggaaaatggccgcttttctggattcatcgactgtggccggctgggtgtggcggacc
gctatcaggaca
tagcgttggctacccgtgatattgctgaagagcttggcggcgaatgggctgaccgcttcctcgtgctt
tacggtatcgcc
gctcccgattcgcagcgcatcgccttctatcgccttcttgacgagttcttctgaattattaacgctta
caatttcctgat
gcggtattttctccttacgcatctgtgcggtatttcacaccgcatacaggtggcacttttcggggaaa
tgtgcgcggaac
ccctatttgtttatttttctaaatacattcaaatatgtatccgctcatgagacaataaccctgataaa
tgcttcaataat
agcacgtgctaaaacttcatttttaatttaaaaggatctaggtgaagatcctttttgataatctcatg
accaaaatccct
taacgtgagttttcgttccactgagcgtcagaccccgtagaaaagatcaaaggatcttcttgagatcc
tttttttctgcg
cgtaatctgctgcttgcaaacaaaaaaaccaccgctaccagcggtggtttgtttgccggatcaagagc
taccaactcttt
ttccgaaggtaactggcttcagcagagcgcagataccaaatactgtccttctagtgtagccgtagtta
ggccaccacttc
aagaactctgtagcaccgcctacatacctcgctctgctaatcctgttaccagtggctgctgccagtgg
cgataagtcgtg
tcttaccgggttggactcaagacgatagttaccggataaggcgcagcggtcgggctgaacggggggtt
cgtgcacacagc
ccagcttggagcgaacgacctacaccgaactgagatacctacagcgtgagctatgagaaagcgccacg
cttcccgaaggg
agaaaggcggacaggtatccggtaagcggcagggtcggaacaggagagcgcacgagggagcttccagg
gggaaacgcctg
gtatctttatagtcctgtcgggtttcgccacctctgacttgagcgtcgatttttgtgatgctcgtcag
gggggcggagcc
tatggaaaaacgccagcaacgcggcctttttacggttcctggccttttgctggccttttgctcacatg
ttctt
