## Supplementary material for "Engineering antigenic breadth against SARS-CoV-2 by pairing divergent RBDs within a single mRNA immunogen": Pain text sequence

GTGCTGAGCTTCGAACTGCTGCATGCCCCTGCCACAGTGTGCGGCCCTAAGAAAAGCACCAATCTCGT
GAAGAACAAAGG
GAGTAGCCAGTGTGTGAACCTGAGAACCAGAACACAGCTGCCTCCAGCCTACACCAACAGCTTTACCA
GAGGCGTGTACT
ACCCCGACAAGGTGTTTAGATCCAGCGTGCTGCACTCTACCCAGGACCTGTTCCTGCCTTTCTTCAGC
AACGTGACCTGG
TTCCACGCCATCCACGTGTCCGGCACCAATGGCACCAAGAGATTCGACAACCCCGTGCTGCCCTTCAA
CGACGGGGTGTA
CTTTGCCAGCACCGAGAAGTCCAACATCATCAGAGGCTGGATCTTCGGCACCACACTGGACAGCAAGA
CCCAGAGCCTGC
TGATCGTGAACAACGCCACCAACGTGGTCATCAAAGTGTGCGAGTTCCAGTTCTGCAACGACCCCTTC
CTGGACGTCTAC
TACCACAAGAACAACAAGAGCTGGATGGAGTTCGGCGTGTACAGCAGCGCCAACAACTGCACCTTCGA
GTACGTGTCCCA
GCCTTTCCTGATGGACCTGGAAGGCAAGCAGGGCAACTTCAAGAACCTGCGCGAGTTCGTGTTTAAGA
ACATCGACGGCT
ACTTCAAGATCTACAGCAAGCACACCCCTATCAACCTCGTGCGGGATCTGCCTCAGGGCTTCTCTGCT
CTGGAACCCCTG
GTGGATCTGCCCATCGGCATCAACATCACCCGGTTTCAGACACTGCTGGCCCTGCACAGAAGCTACCT
GACACCTGGCGA
TAGCAGCAGCGGATGGACAGCTGGTGCCGCCGCTTACTATGTGGGCTACCTGCAGCCTAGAACCTTCC
TGCTGAAGTACA
ACGAGAACGGCACCATCACCGACGCCGTGGATGGGTGGTACATCTGGCTGGGCTTTATCGCCGGACTG
ATTGCCATCGTG
ATGGTCACAATCATGCTGTGTTGCATGACCAGCTGCTGTAGCTGCCTGAAGGGCTGTTGTAGCTGTGG
CAGCTGCTGCAA
GTTCGACTGATGAGAATTCgctggagcctcggtggccatgcttcttgccccttgggcctccccccagc
ccctcctcccct
tcctgcacccgtacccccgtggtctttgaataaagtctgagtgggcggcagcctgtgtgtgcctgagt
tttttccctcag
caaacgtgccaggcatgggcgtggacagcagctgggacacacatggctagaacctctctgcagctgga
tagggtaggaaa
aggcaggggGCggccgctcgagtctagagggcccgtttaaacccgctgatcagcctcgactgtgcctt
ctagttgccagc
catctgttgtttgcccctcccccgtgccttccttgaccctggaaggtgccactcccactgtcctttcc
taataaaatgag
gaaattgcatcgcattgtctgagtaggtgtcattctattctggggggtggggtggggcaggacagcaa
gggggaggattg
ggaagacaatagcaggcatgctggggatgcggtgggctctatggcttctactgggcggttttatggac
agcaagcgaacc
ggaattgccagctggggcgccctctggtaaggttgggaagccctgcaaagtaaactggatggctttct
cgccgccaagga
tctgatggcgcaggggatcaagctctgatcaagagacaggatgaggatcgtttcgcatgattgaacaa
gatggattgcac
gcaggttctccggccgcttgggtggagaggctattcggctatgactgggcacaacagacaatcggctg
ctctgatgccgc
cgtgttccggctgtcagcgcaggggcgcccggttctttttgtcaagaccgacctgtccggtgccctga
atgaactgcaag
acgaggcagcgcggctatcgtggctggccacgacgggcgttccttgcgcagctgtgctcgacgttgtc
actgaagcggga
agggactggctgctattgggcgaagtgccggggcaggatctcctgtcatctcaccttgctcctgccga
gaaagtatccat
catggctgatgcaatgcggcggctgcatacgcttgatccggctacctgcccattcgaccaccaagcga
aacatcgcatcg

agcgagcacgtactcggatggaagccggtcttgtcgatcaggatgatctggacgaagagcatcagggg
ctcgcgccagcc
gaactgttcgccaggctcaaggcgagcatgcccgacggcgaggatctcgtcgtgacccatggcgatgc
ctgcttgccgaa
tatcatggtggaaaatggccgcttttctggattcatcgactgtggccggctgggtgtggcggaccgct
atcaggacatag
cgttggctacccgtgatattgctgaagagcttggcggcgaatgggctgaccgcttcctcgtgctttac
ggtatcgccgct
cccgattcgcagcgcatcgccttctatcgccttcttgacgagttcttctgaattattaacgcttacaa
tttcctgatgcg
gtattttctccttacgcatctgtgcggtatttcacaccgcatacaggtggcacttttcggggaaatgt
gcgcggaacccc
tatttgtttatttttctaaatacattcaaatatgtatccgctcatgagacaataaccctgataaatgc
ttcaataatagc
acgtgctaaaacttcatttttaatttaaaaggatctaggtgaagatcctttttgataatctcatgacc
aaaatcccttaa
cgtgagttttcgttccactgagcgtcagaccccgtagaaaagatcaaaggatcttcttgagatccttt
ttttctgcgcgt
aatctgctgcttgcaaacaaaaaaaccaccgctaccagcggtggtttgtttgccggatcaagagctac
caactctttttc
cgaaggtaactggcttcagcagagcgcagataccaaatactgtccttctagtgtagccgtagttaggc
caccacttcaag
aactctgtagcaccgcctacatacctcgctctgctaatcctgttaccagtggctgctgccagtggcga
taagtcgtgtct
taccgggttggactcaagacgatagttaccggataaggcgcagcggtcgggctgaacggggggttcgt
gcacacagccca
gcttggagcgaacgacctacaccgaactgagatacctacagcgtgagctatgagaaagcgccacgctt
cccgaagggaga
aaggcggacaggtatccggtaagcggcagggtcggaacaggagagcgcacgagggagcttccagggg
gaaacgcctggta
tctttatagtcctgtcgggtttcgccacctctgacttgagcgtcgatttttgtgatgctcgtcagggg
ggcggagcctat
ggaaaaacgccagcaacgcggcctttttacggttcctggccttttgctggccttttgctcacatgttc
tt
