## Supplementary Figure 1 for "Engineering antigenic breadth against SARS-CoV-2 by pairing divergent RBDs within a single mRNA immunogen"

### Supplementary Material

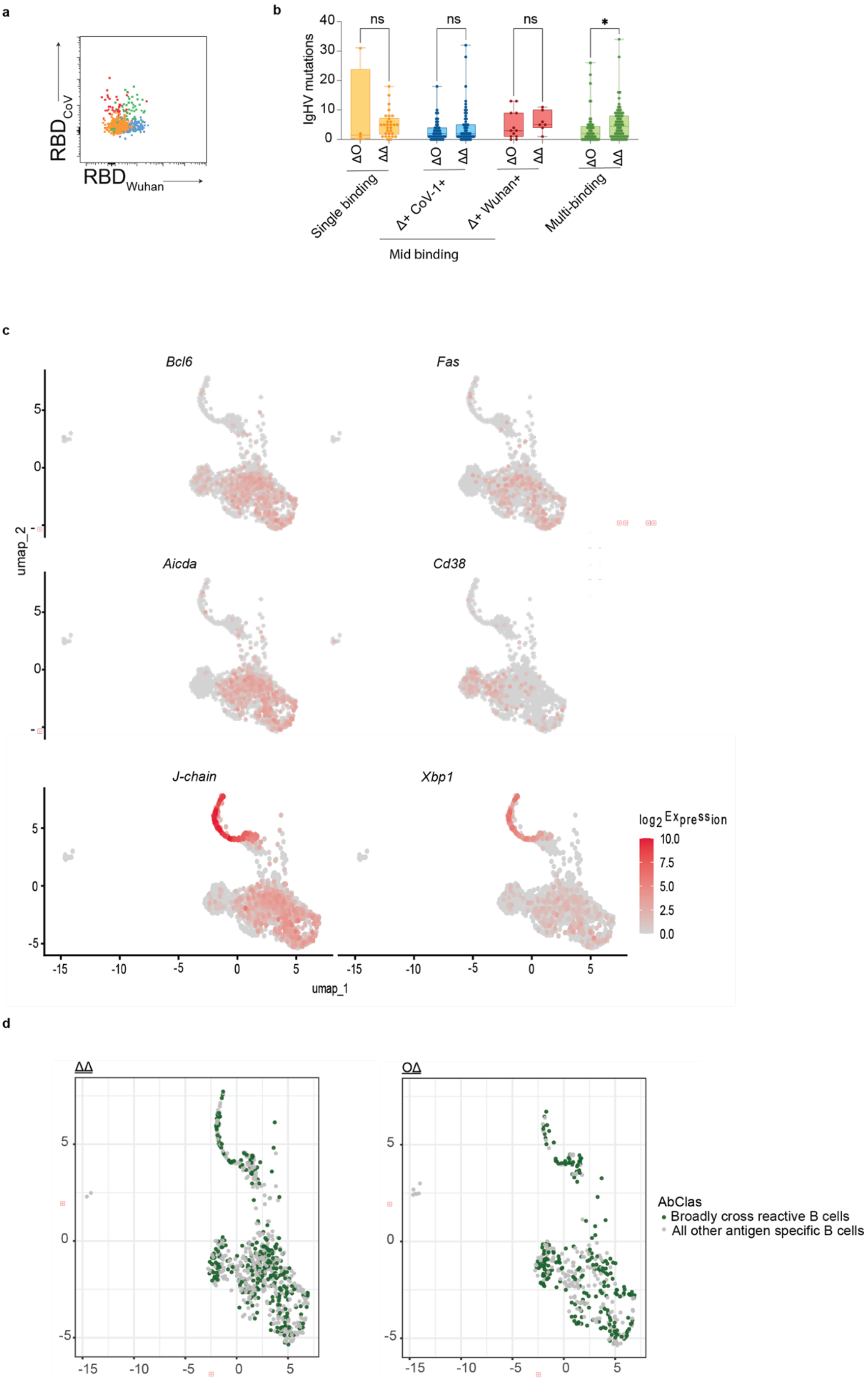

**Supplementary Figure 1: a,** Efficacy of surrogate staining confirmed by flow cytometry, using fluorescently tagged antibodies in place of oligotagged antibodies. **b,** level of SHM in cells from different tiers of polyreactivity from mice with immunised with the matched (Δ-Δ) or divergent (O-Δ) tRBD-NTD. **c,** expression of GC cell markers, memory markers and plasmablast markers among clusters and of antigen specific B cells. **d,** clustering of multi-binding cells (green) as compared to all over RBD binding B cells (grey) among B cells from mice immunised with the divergent or matched (Δ-Δ) vaccine.
